# Multiscale modelling of invasive species control: the cane toad in Australia

**DOI:** 10.1101/2022.12.07.518288

**Authors:** Anh D. Pham, Christopher M. Baker, Nicholas Geard

**Affiliations:** School of Computing and Information Systems, The University of Melbourne, Parkville, Victoria 3010, Australia; School of Mathematics and Statistics, The University of Melbourne, Parkville, Victoria 3010, Australia

**Author notes:** College of Engineering and Computer Science, VinUniversity, Hanoi, Vietnam (Anh D. Pham).

**Keywords:** agent-based model, *Bufo marinus*, control, invasive species, multiscale model, simulation, spread

## Abstract

1. The cane toad (*Bufo marinus*) is an invasive species in Australia that has a negative impact on native species. Control methods such as trapping, fencing, and water exclusion have been devised to contain the spread of cane toads and reduce their ecological impact. However, implementing these interventions is expensive, and estimating the likely impact of a proposed intervention on spread at a large spatial scale, comprising one or more control methods, is challenging due to the lack of large-scale data and the computational cost of modelling a large number of toads.
2. To address this challenge, we developed a multiscale model which uses individual-level data on cane toad behaviour to estimate the impact of trapping, fencing, and water exclusion when applied at scale in the Pilbara region in north-western Australia. Compared to previous work, our model allows us to explore more complex combinations and tradeoffs of control methods by utilising data sources at different scales.
3. Our results suggest that exclusion of toads from water points is the most effective method for containing the spread of cane toads, and that trapping and fencing alone are unlikely to be sufficient. However, trapping and fencing are still useful supplementary measures in scenarios where exclusion cannot be broadly applied to a large number of water points.
4. *Synthesis and applications*. Our analyses highlight the importance of limiting access to sheltering and breeding sites in invasive species control. Furthermore, this study illustrates the value of multiscale computational models for exploring scenarios where parameters and calibration data are available at the scale of individuals and small groups, but management questions are framed at a much larger scale.

## Introduction

Invasive species can cause great harm to the local ecosystems, and controlling their spread can be challenging. One example is that of the Cane toad (*Bufo marinus*), which was introduced to Queensland, Australia in 1935 in an effort to control the pest cane beetle (Easteal 1981). Since its introduction, the toad has spread through several states in accelerating rates Shine et al. (2021), Phillips et al. (2010) and reaches billions in population (Tingley et al. 2017), causing significant damage to local ecosystems. As of 2022, cane toads are present in the Kimberley region in Western Australia and are advancing towards the Pilbara region (Department of Biodiversity, Conservation and Attractions, Western Australia 2021, 2022) (Figure S6, Supporting Information).

Given their potential impact, a variety of methods have been proposed to manage cane toads, including manual collection, trapping and fencing. Manual collection by local communities and toad traps (Peacock 2007) have proven useful to eradicate some satellite populations (Wingate 2011). Traps with both insect-attracting light and acoustic lure – recorded advertisement calls (Yeager et al. 2014, Muller & Schwarzkopf 2017) – have been found to be more labour-effective than manual collection. Crossland et al. (2012) developed another type of trap that targets the tadpole phase of toad metamorphosis, which have proven effective at capturing a large number of tadpoles and reducing the presence of toads afterwards (Crossland et al. 2012, McCann et al. 2019). Besides traps, previous studies have explored the use of physical fences to keep toads out of conservation areas (Brook et al. 2011). In more arid regions, cane toads rely on natural and artificial water points to survive during dry season (Florance et al. 2011), and removing their access to such water points can be an effective control method (Letnic et al. 2015, Gregg et al. 2019). Strategies to control the spread of invasive species often involve significant resources and long-term commitment. It is therefore important to optimise their design before they commence. Mathematical and computational models are a cost-effective approach to evaluating and comparing a wide range of strategies while accounting for uncertainties in environmental conditions.

Models have previously been used to estimate the potential range of cane toads based on their physiological constraints (Kearney et al. 2008), climatic conditions of their home range (Sutherst et al. 1996) and environmental and anthropogenic factors of their current range in Australia (Urban et al. 2007, Doody et al. 2018). Several studies also employed models to investigate the enhanced dispersal capability of toads at the invasion front (Burton et al. 2010, Bouin et al. 2018, Alex Perkins et al. 2013). Models built to evaluate control strategies are less common. Florance et al. (2011) employed modelling to investigate the toad’s reliance on water points in arid regions and explore the effect of blocking the toad’s access to such water points. Later studies have followed up by exploring the potential of a waterless barrier, created by excluding toads from artificial water points in the Kimberley-Pilbara corridor to prevent toads from spreading further into the Pilbara (Tingley et al. 2012, Southwell et al. 2017) (from now on referred to as “exclusion”). Notably, recent models of toad spread share a similar approach: predicting large-scale spread from individual-level movement data through an intermediate step. For example, Tingley et al. (2012) used movement data to generate a 2D dispersal kernel and used this kernel to model spread across the large Kimberley-Pilbara region.

Despite the wealth of possible control methods, progress has been slow in stopping the cane toad invasion (Tingley et al. 2017). Although some control methods such as fencing and trapping have been proven to be effective on a small scale (Muller & Schwarzkopf 2018) (less than 1 square kilometre), little has been done to model the impact and cost of deploying them on a larger scale. While some studies have confirmed the large-scale effectiveness of exclusion (Tingley et al. 2012, Southwell et al. 2017), this strategy requires significant funding and commitment from the state government, relevant departments and private landowners, and as such need further investigation and risk assessment before it can be realised. Finally, to our best knowledge there have been no attempts to estimate the impact of combining multiple control methods on a large scale (e.g. an area spanning thousands of square kilometres).

In this study, we modelled the spread of cane toads in the Kimberley-Pilbara region of Australia in order to estimate the likely impact of a combination of trapping, corridor-fencing, and exclusion.

Specifically, we seek to answer the question: What is the large-scale impact of trapping and corridor-fencing on the spread of cane toads in the Kimberley-Pilbara corridor, especially when combined with exclusion? To answer this question, we employed a multiscale model, similar to the approach taken in (Tingley et al. 2012); however, we used colonisation probability between water points instead of a dispersal kernel to model population-level spread. This multiscale model allows us to utilise data at individual level, which are more readily available, to effectively represent a large number of agents on a relatively large spatiotemporal scale. From simulation results of alternative control scenarios and parameters, our model suggests that toads can only be stopped by removing their access to water points, and trapping and corridor-fencing can still supplement this strategy and reduce the impact of the uncertainty in parameters.

The rest of the paper is structured as follows. In Materials and methods, we describe our multi-scale model and detail the input data used to parameterise it. In Results, we present the simulations, conducted with the model, to estimate the impact of control methods and the corresponding results. In Discussion, we discuss how our results demonstrate that trapping and fencing alone are unlikely to stop toads, as well as the implications of our study to cane toad control and invasive species modelling.

## Materials and Methods

We constructed a multiscale model comprised of two connected agent-based models. Each model included a representation of space, but at very different scales. The first (macroscale) model represents population-level spread in the Kimberley-Pilbara corridor. The second (microscale) model of toads’ individual behaviour in relatively small areas.

### The macroscale model of spread and control

The macroscale model is configured to represent the Kimberly-Pilbara region and allows us to make estimates regarding the spread of toads in the region and impact of control on this spread by utilising the output of the microscale model. In the macroscale model, water points in the Kimberley-Pilbara corridor are modelled as a connected network of nodes, and the large-scale spread of cane toads through this region as transmission through the water point network. The links between water points determine the probability of toads spreading from one water point to another. We estimated this probability using the microscale model and it depends on variables such as the distance between them, the capacity of the colonised water point, and the climate in each subregion. Control methods further modify this probability, reducing it to different extents. In each simulation run, toads spread through the corridor from a number of pre-colonised water points in the North-Eastern end (Figure 1). Each simulation ends when toads reach the South-Western end of the corridor or after 200 years.

**Figure 1:**
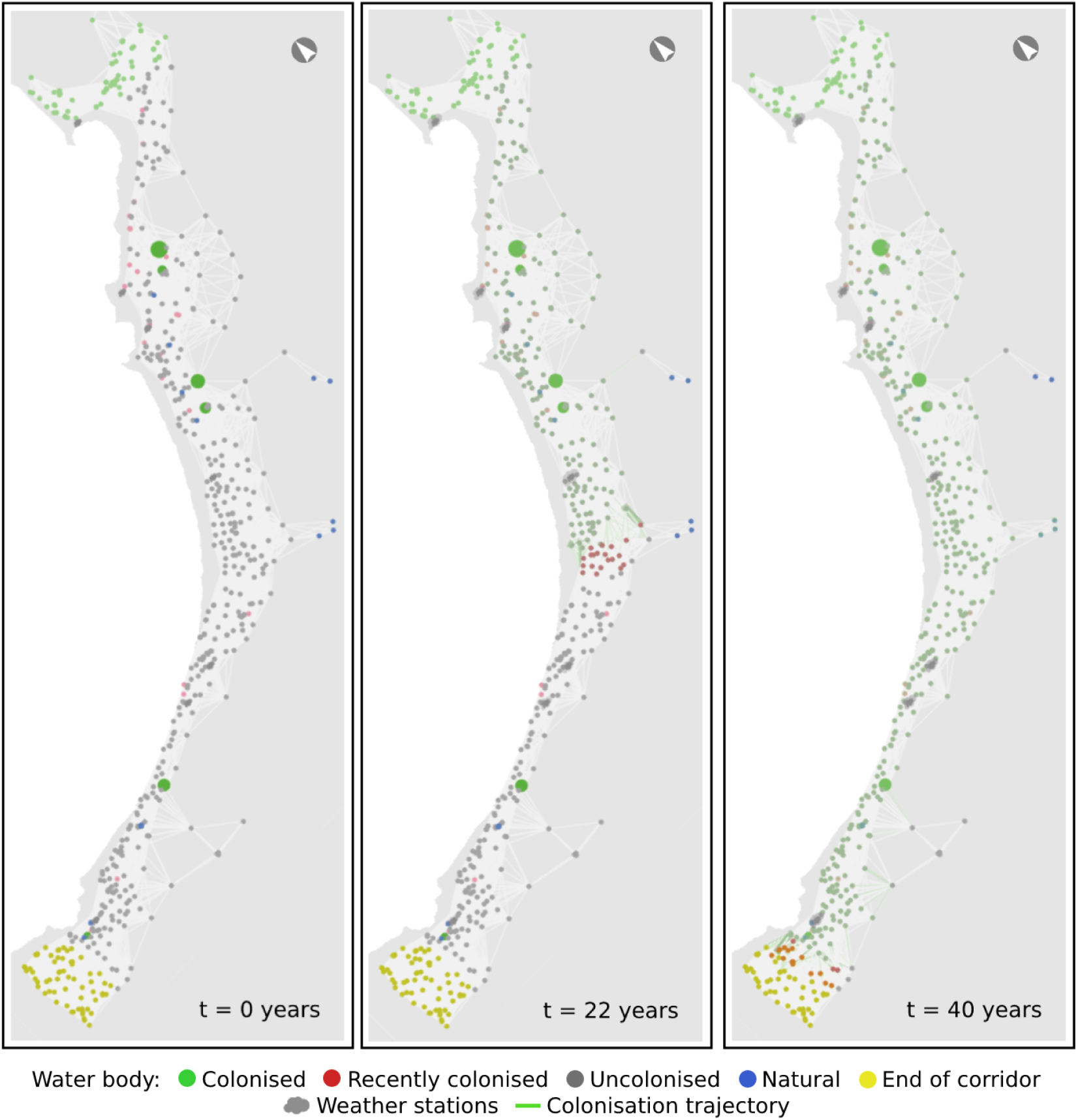
Output from a typical simulation run showing toads spreading along the Kimberley-Pilbara corridor through time. At the beginning of the simulation (left panel; t = 0 years) toads are concentrated in the Northern water points. After 22 years (centre panel), toads have reached the centre of the corridor, and by 40 years (right panel), toads have reached the Southwestern end of the corridor.

The distribution of water points in the Kimberley-Pilbara corridor affects the spread dynamics, as well as the effectiveness of control strategies, and thus is crucial for the macroscale model. We use data from the most recent model of cane toads in this region (Southwell et al. 2017). The data were aggregated from several sources, including Geoscience Australia (Geo n.d.), Department of Water, WA and Department of Agriculture and Food, WA, and verified by land managers in this region. The dataset contains 566 water points, each with location (in coordinates) and type (natural, dam, tank, irrigation, dwelling, etc.). Water points of the type “irrigation area” also have size measurements in the form of area and perimeter. In addition to water point distribution, rainfall pattern directly impacts how fast and how far toads can spread due to their reliance on rainfall during wet seasons. Variations in rainfall between subregions of the corridor are modelled and parameterised using weather records of weather stations situated along the corridor (*Bureau of* Meteorology n.d.).

In the macroscale model, we model three control methods: trapping, corridor-fencing, and exclusion. A control location is defined as an area where control methods will be concentrated. 17 control locations are generated uniformly along the corridor, from location 0 at the South-Western end to location 16 at the North-Eastern end (Figure S4, Supporting Information). In each scenario with control methods, one control location is chosen. Exclusion and trapping are employed at a number of water points closest to the control location’s centre. Corridor-fencing involves a corridor-wide fence positioned at the beginning of the control location and perpendicular to the length of the corridor (Figure S5, Supporting Information).

Corridor-fencing affects the probability that toads, from water points on one side of the fence, colonise water points on the other side. Trapping affects the probability that the controlled water points are colonised. Opposed to corridor-fencing and trapping, which are represented in the microscale model and reduce colonisation probability between water points, the exclusion control method is represented in the macroscale model to replicate previous studies (Tingley et al. 2012, Southwell et al. 2017). Exclusion restricts access to water points where it is applied, but exclusion at each water point has a 5% probability to fail and allow toads to colonise the corresponding water point. We assume that exclusion failures are detected and repaired after two years.

Further details on the macroscale model can be found in Section S2, Supporting Information.

### The microscale model of toad movement

The microscale model is a spatial abstraction that allows us to input fine-grained movement rules for toads and observe the probability of colonisation of water points under different initial conditions. In the microscale model, each toad is modelled as an agent that moves away from a colonised water point and across a relatively small landscape (under 200 km^2^), potentially colonising new water points (Figure 2). It simulates scenarios where individual-level behaviour matters, such as at the invasion front and when environmental conditions permit the dispersal of toads. The output of the model - the probability of colonisation between water points - is later used to parameterise the large-scale model.

**Figure 2:**
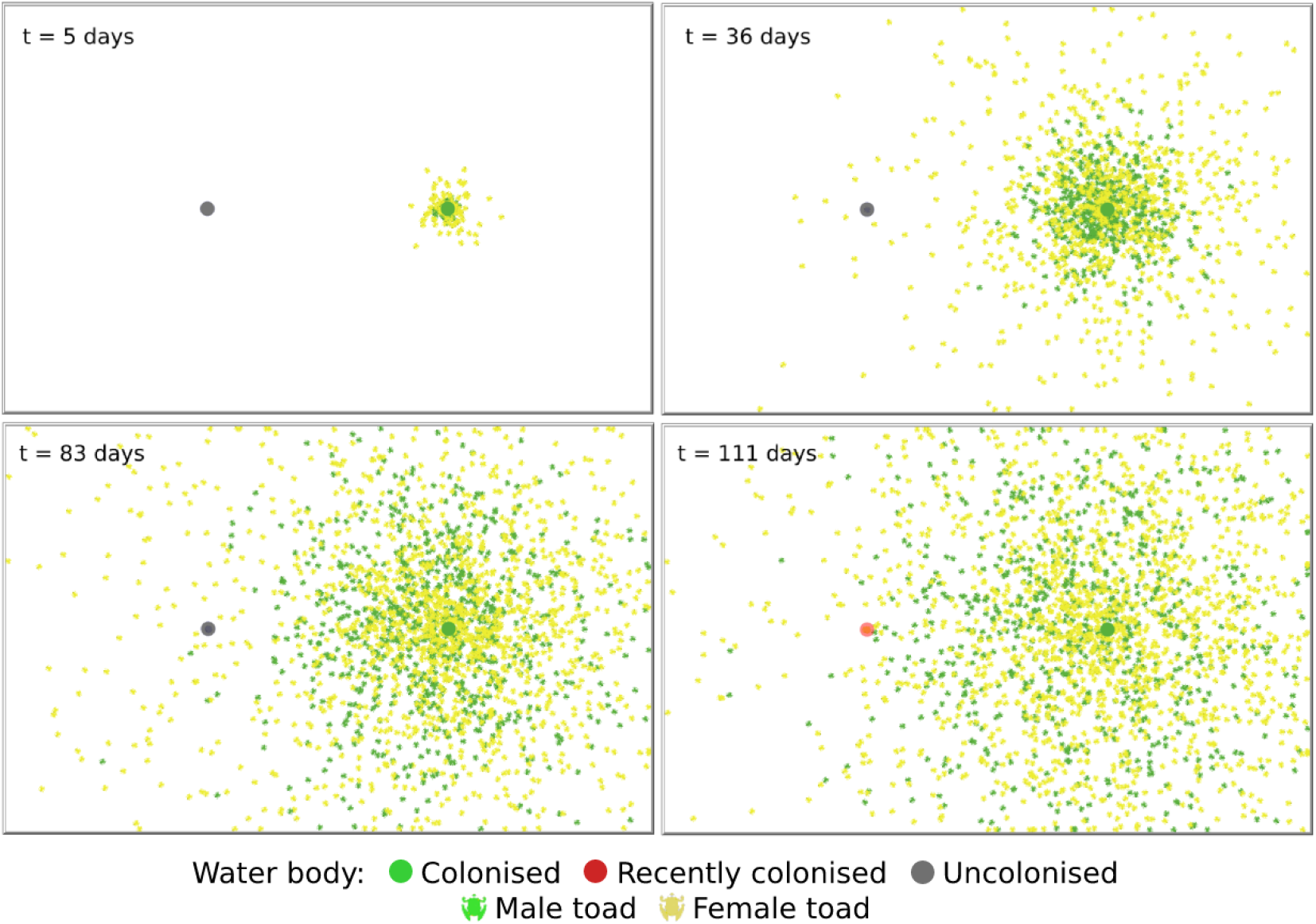
Toads disperse and colonise a water point 6 kilometres away in a simulation run. At the beginning of the simulation (top left panel; t = 5 days) toads start to leave the source water point. After 36 days (top right panel), due to dispersing faster female toads have reached the uncolonised water point first. By 111 days (bottom right panel), male toads have arrived, which results in the colonisation of the water point.

Movement rate is a key driver of cane toad spread, and was drawn from (Phillips et al. 2007) in which the movement of many cane toads were tracked and recorded in the wet-dry tropics of Northern Territory, a region with rainfall pattern similar to the modelled area. We define active days as the days toads can move freely during wet seasons, and assume that toads move at a constant rate during active days. We assume toads emerge from a colonised water point at a rate of hundreds of thousands per generation (i.e. around 10^3^ per day) and can detect water from 100m away - Tingley et al. (2012) made similar assumptions based on expert opinion and calibrating to spread data. Other ecological characteristics of toads are also essential in constructing the model, such as sex ratio, time to reach adulthood, and the fact that juvenile toads do not venture far from their water point (Lever 2001, Hayes et al. 2006, Can n.d.).

We model traps as locations in space that have a probability of capturing each toad within a specified radius. To our knowledge, there have been no previous models of cane toad traps. However, in a previous individual-based model of invasive species control (Hradsky et al. 2019), an approach similar to ours was employed – individuals with a territory overlapping a bait station are at risk of dying from that station. Trap radius and probability of capture were parameterised using data from field experiments (Muller et al. 2016, Muller & Schwarzkopf 2017, 2018), which reported 3% daily capture rate and 120-meter range. We assume that traps are arranged uniformly in a grid - a common practice in management - and that they are placed 100m away from each other in the area of 0.04 square kilometres around the water point.

We model a fence as contiguous locations in space that cannot be crossed by moving cane toads. Each section of a fence has a daily probability of becoming damaged; for example, by natural phenomena, wildlife and human activity (Letnic et al. 2015). Toads can then move freely through a damaged fence section until it is repaired. In the absence of more detailed data on fence robustness, we assume that a fence section has a 5% probability of being damaged each day. All damaged fence sections are repaired at the beginning of each week.

Further details on the microscale model can be found in Section S1, Supporting Information.

### Simulations and Results

We used the multiscale model described above to quantify how the rate of spread and the probability that toads will colonise the studied region vary with the application of control methods. In addition, we run sensitivity analyses on important assumed parameters to explore how uncertainty affects such predictions.

### Estimating toad spread between water points - with and without control

This set of simulations estimates the spread of cane toads between two water points when control methods are not present and when control methods are present. We made certain assumptions regarding unknown control parameters, and ran simulations using the parameters detailed in Table S2 (Supporting Information). There are 4 scenarios in total: no control; fencing-only; trapping-only and fencing-trapping; and for each scenario we explore a range of distance (6 to 24km) and active days (40 to 160 days). We ran 1000 simulations for each parameter combination and afterwards the results are aggregated to obtain the colonisation probability.

We found that the spread of cane toads between water points is strongly dependent on distance and active days. Specifically, at 160 active days, colonisation probability is relatively high (more than 50%) regardless of capacity as long as the distance is less than 8 km (Figure 3a). As distance increases, colonisation probability starts to drop off quickly; at a distance above 16 km, colonisation probability is often less than 5%. Decreasing active days has a similarly significant effect – toads can only colonise a water point more than 14 km away at 160 active days. At 40 active days (minimum value), colonisation probability is extremely low save for the shortest distance (6 km). In other words, the distance toads can spread is strongly limited by the number of active days.

**Figure 3:**
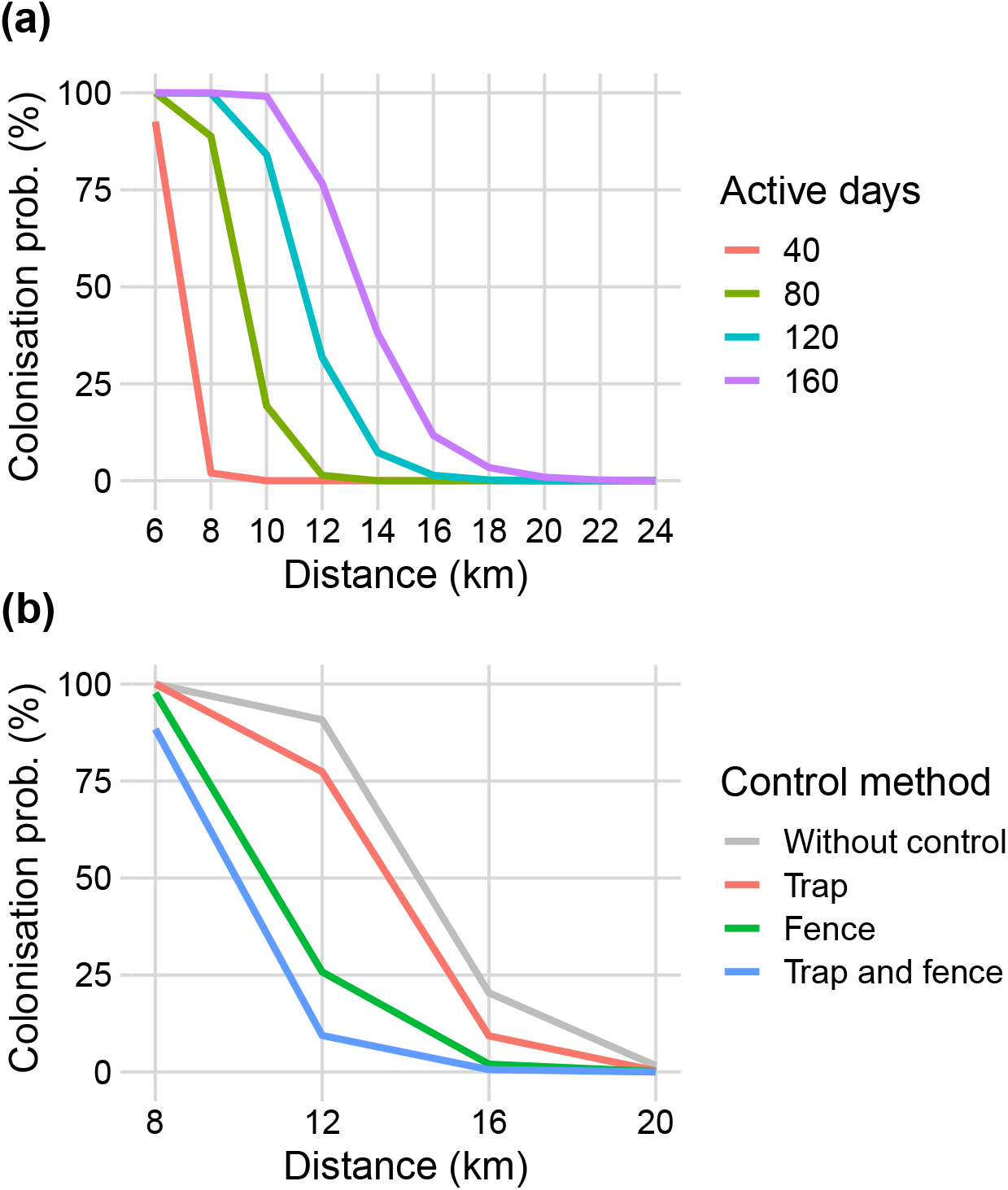
Relationship between number of active days and colonisation probability at a given distance. The impact of fencing and trapping is only noticeable at medium distance.

Control methods have differing impact on toads’ ability to spread. Overall, the results show that, under our assumptions in trap density and fence robustness, trapping has a lesser impact compared to fencing. The impact of trapping is more pronounced when both methods are deployed together, in which case colonisation probability is significantly reduced. Moreover, both control methods only have noticeable impact at medium distance (from 8km to 16km).

### Estimating toad spread in Kimberley-Pilbara corridor, with and without control

Here we use the macroscale model to estimate how long cane toads will take to spread through the Kimberley-Pilbara region. Parameters take default values described in Table S4 (Supplementary File) with and without control methods. We ran 2000 simulations and observed the time it takes for the region to be colonised. The simulation results are aggregated to produce the mean and the range of outcomes. The results provide a baseline to compare against those of later simulations.

In addition to no-control scenario, we simulate five controlled scenarios, each involving either a single control method or a combination of exclusion and other control methods (corridor-fencing; trapping; exclusion; exclusion and corridor-fencing; and exclusion and trapping). The control location is varied between 17 pre-generated control locations running along the corridor and the number of controlled water points ranges from 4 to 14. We run 1000 simulations for each combination and observe whether the region is colonised in 200 years. The results are aggregated to produce the probability of the region being colonised.

According to our model, without any intervention, the Kimberley-Pilbara corridor will be colonised in 47 to 64 years. The mean estimated time is 52.6 years and the standard deviation is 2.1 years.

Between the three control methods, only exclusion is likely to stop the invasion completely. The best control locations at the North-Eastern end, the middle and South-Western end of the corridor are locations 3, 10 and 15 respectively - other control locations in each respective section perform similarly or worse in terms of preventing colonisation. When only exclusion is deployed, at some locations (“No fence” scenario at location 15 in Figure 5), it takes as few as 10 controlled water points to achieve a region colonisation probability lower than 5%. Controlling more water points brings this probability down further while also making more locations viable (Such as location 3).

When used to supplement exclusion, corridor-fencing has a more noticeable impact (Figure 5). In certain scenarios, adding corridor-fencing is more effective at reducing the probability of the region being colonised compared to deploying exclusion at several more water points (control location 10), and thus potentially allows securing the region (colonisation probability less than 0.05) with less water points controlled (control location 3). Trapping follows a comparable pattern, although as shown previously its impact is likely much weaker than that of fencing under our assumptions.

### Exploring sensitivity to capacity of water points and wet climate

Here we explore the sensitivity of the model with regard to the mean capacity of water points. This parameter is unknown but potentially impactful, as it can enable toads to spread between water points further apart than predicted in the microscale level. We re-run the scenarios to estimate toad spread with and without control methods, while assuming different values for the mean capacity of water points to explore the effect on spread and evaluate the robustness of control strategies. Specifically, mean capacity is set at 240 and 3840 colonisers per day (i.e. default value, 960, divided and multiplied by 4).

According to our model, increasing the mean capacity of water points by a factor of 4 reduces the expected colonisation time drastically. Specifically, when mean capacity of water points is assumed to be 3840 toads per day, according to the results the region is colonised in 36 to 46 years, and in 41.3 years on average – 12.3 years earlier. For the lower assumption of capacity (240 toads per day), this range is instead 64 to 98 years, with the average time being 77.5 years – 24.9 years later.

A larger mean water point capacity allows toads to spread further than before and renders many control strategies ineffective. Control location 15 is the most robust, with the region still protected from colonisation (colonisation probability less than 5%) even with the highest assumption of capacity, given that 14 water points are controlled. Finally, in scenarios where exclusion is employed at more water points to compensate for higher capacity (up to 20 from 14 water points), some control locations become viable again (location 3 in Figure 5), while a few locations remain ineffective regardless (location 10). On the other hand, if our assumption turns out to be an overestimate and capacity is indeed lower, more locations can contain the spread of toads through the region (such as location 10). Strategies that combine exclusion and another control method are also less effective in high-capacity scenarios.

Climate is another aspect of the model subject to uncertainty. As shown previously, the number of active days in a wet season influences colonisation probability between water points and can have a similar impact on large-scale spread. Although the number of active days is informed by data, there is still uncertainty due to future climate change. We re-ran the previous simulations to estimate toad spread with and without control methods, this time reducing the interval between wet years and increasing their intensity to explore the effect on spread and evaluate the robustness of control strategies. Specifically, the interval between wet years is set at 5 years instead of the default value of 10 years, and the number of extra active days during wet years is set at 28 days instead of the default value of 14 days.

Our model predicts that that wetter climate is likely to speed up the spread of cane toads to a small extent. When wet years occur twice as often and with stronger impact, toads can colonise the corridor in 43 to 46 years, with the expected colonisation time being 48.8 years – 3.8 years quicker on average than under current assumptions (Figure 4). Similarly, wetter climate slightly reduces the effectiveness of control strategies. Control locations 3 and 15 remain effective even with exclusion as the only control method deployed. Combining exclusion and corridor-fencing results in slightly more robustness to changes in climate, as evidenced by location 10 (Figure S7, Supporting Information).

**Figure 4:**
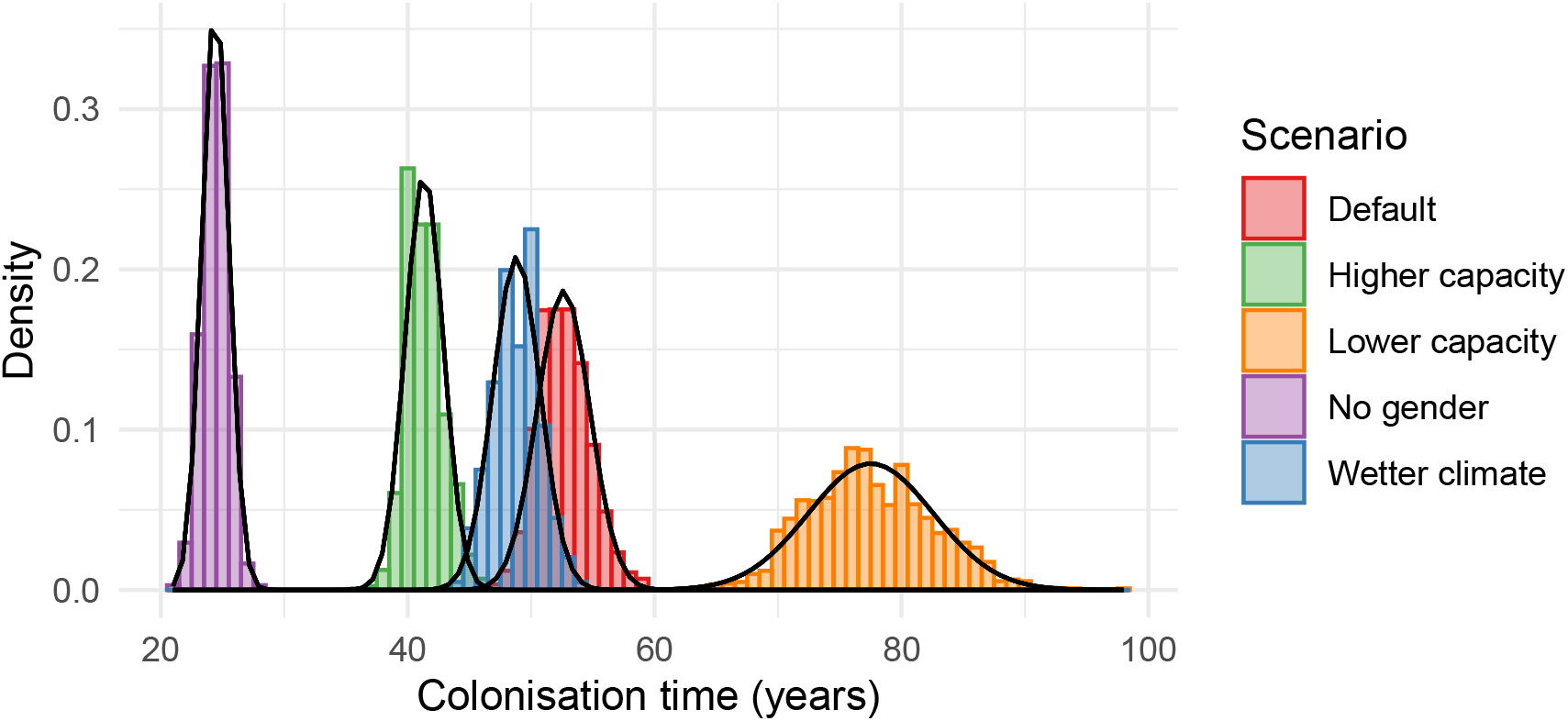
Colonisation time distribution of all no-control scenarios.

**Figure 5:**
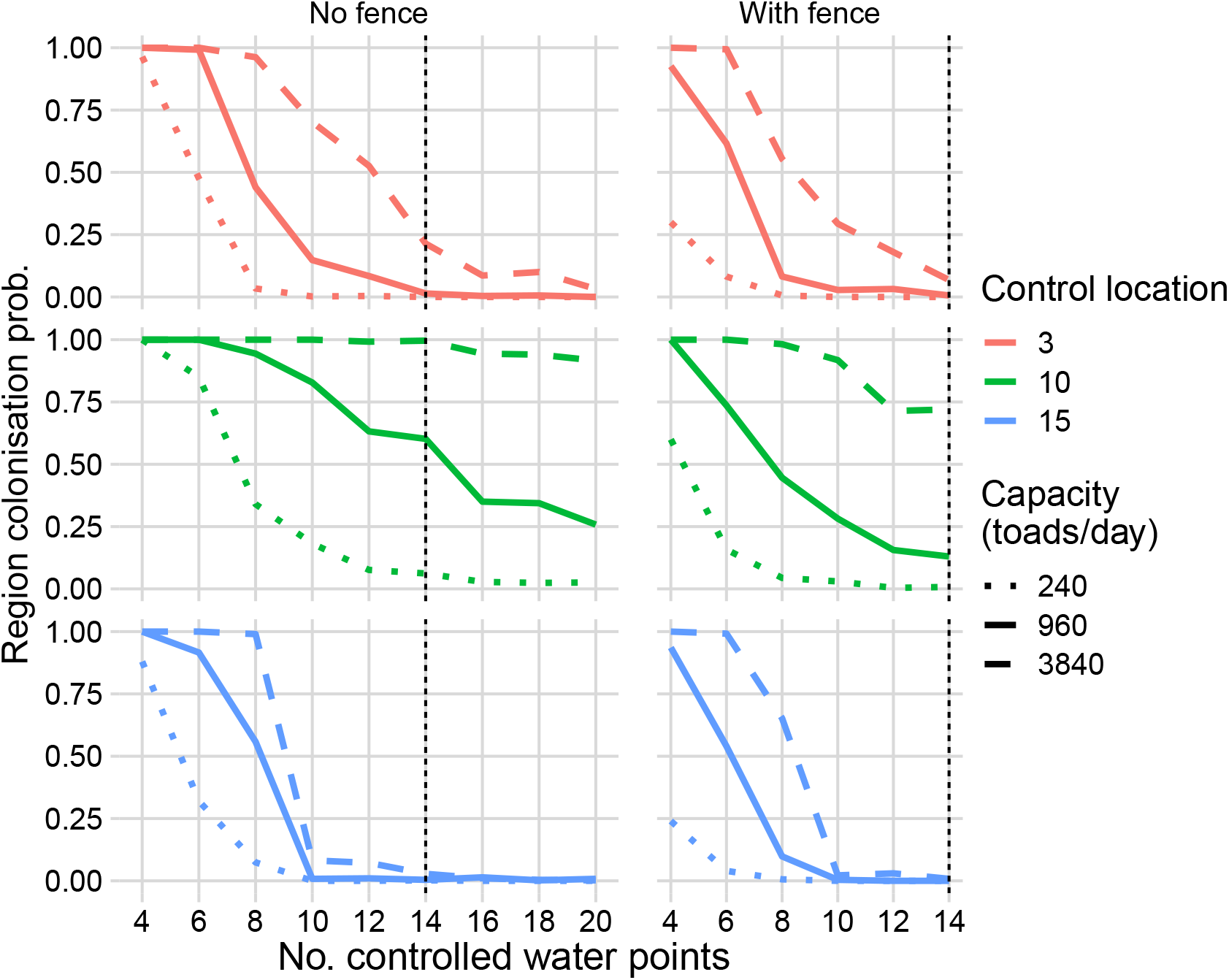
Higher capacity can lead to drastic changes in outcome. The colored lines represent colonisation probability under a combination of exclusion and corridor-fencing; solid lines are the results under default assumptions, dotted lines and dashed lines are the results under lower and higher assumption of capacity, respectively.

## Discussion

We investigated the large-scale impact of trapping and corridor-fencing on the spread of cane toads in the Kimberley-Pilbara corridor, when combined with exclusion. The results of our model suggest that without management, cane toads will colonise the Kimberley-Pilbara region within 47-64 years. Under the assumptions made in this study, trapping and corridor-fencing are unlikely to halt the spread of cane toads when used in isolation, and to do so will likely require exclusion - restricting their access to a number of water points. The best locations to stop the spread are at the ends of the corridor. Despite having little effect when used in isolation, trapping and corridor-fencing can increase the effectiveness of exclusion. Finally, although a wetter climate and uncertainty in water point capacity might result in faster spread and reduced protection from control strategies, they can be accounted for by employing exclusion at more water points or supplementing an exclusion strategy with trapping or a corridor-fencing. The exact amount of compensation needed (i.e. number of extra water points to control or intensity of secondary control methods) can vary depending on the location and the anticipated scenarios.

Our study confirms that trapping and corridor-fencing are unlikely to be effective at stopping toads from crossing the Kimberley-Pilbara corridor. The inability of corridor-fencing and trapping to stop toads at large scale can be attributed to the ineffectiveness of fencing and trapping at short distances and the distribution of water points in the Kimberley-Pilbara corridor, with sections of the corridor consistently within 5 km of each other. For both methods to have some impact, colonisation trajectories need to be longer (i.e. sparser distribution of water points), and there must be frequent monitoring to de-colonise water points. This is evidenced by their increased effectiveness when used in a “mostly waterless zone” created by exclusion, as this results in the toad’s increased reliance on medium- and long-distance spread to “hop” through this zone and colonise the entire region. In addition, trapping and fencing might prove more effective under other conditions, such as more effective traps, more robust fence material, or more frequent fence maintenance. In many cases, the impact of adding corridor-fencing or trapping to an exclusion-only strategy is comparable to the impact of employing exclusion at more water points. However, in practice some water points cannot be controlled for different reasons (Southwell et al. 2017) – such as the nature of the water point or the cooperation of landowners – and in such cases, supplementing exclusion with another control method such as trapping or corridor-fencing remains a reasonable alternative.

Our approach is similar to that employed by Tingley et al. (2012); however, we used colonisation probability between water points instead of a dispersal kernel to model population-level spread. This approach allows us to model the impact of control methods more directly. Compared to previous estimates (Tingley et al. 2012, Southwell et al. 2017), our model estimates a slower rate of spread through the region. This difference likely results from the inclusion of gender in our model and the large gap in movement rate between male and female cane toads. We removed gender from the model (i.e. only requiring two toads of any gender present at a water point to colonise it, similar to previous models) and observed that toads were able to spread much faster (Figure 4), leading to estimates that were more comparable to those made by previous studies. Specifically, when gender is not modelled, toads would take 24.4 years to colonise the region compared to previous predictions of 25 years (Tingley et al. 2012) and 20.3 years (Southwell et al. 2017). Although in our model, strategies that employ exclusion require managing less water points – most likely due to the slower spread – the optimal locations remain roughly similar, with best locations being at two ends of the corridor (locations 3 and 15 in this study).

Modelling cane toad control in the Kimberley-Pilbara region posed a methodological challenge: the need to model toad movement and control at an individual scale where data are more available while also making predictions regarding the impact of control strategies, which span a large spatiotemporal scale. We addressed these challenges in methodology by constructing a multiscale agent-based model consisting of a microscale model, which makes use of individual-level data, and a macroscale model, which represents spread and control at a larger scale. Utilising all available data at both scales allows us to incorporate more control methods and make predictions that are more grounded - evidenced by how modelling gender changes the predicted outcomes significantly. In the future, as more data on toad movement become available, the model can be easily re-parameterised to update its large-scale predictions. Furthermore, by separating and integrating the two models efficiently – such as only representing toads in small spatial domains and reusing the microscale simulation results across the large landscape – we significantly reduce the computational cost compared to that of a single-scale model. Finally, this approach facilitates the analysis of the microscale model in isolation, as its outcomes can be easily extracted.

Beyond cane toads, the model is also suitable to represent the spread and control of other invasive species with similar characteristics, such as a strongly seasonal dispersal cycle and a heavy reliance on habitat patches. The multiscale design even facilitates the adaptation of the model to a different invasive species. For example, the spread of another invasive species might rely on similar macroscale processes such as climate conditions and distribution of habitat patches, despite the individuals of that species having different characteristics and behaviours from cane toads. In that case, a new microscale model can be constructed, and the simulation results of this new model can still be used as parameters in the existing macroscale model to make predictions about the spread and control of that species in a real landscape.

There remain several limitations to our studies. In terms of model parameters, we had to make similar assumptions to previous models regarding the rate of toads leaving water points in search of new habitats. This prevalent uncertainty in cane toad models highlights the need for future studies to narrow down the range of emigration rate of cane toads from water points to facilitate better estimates. Moreover, long-distance dispersal - incidents where toads hitch-hike on vehicles to travel greater distances - can also affect the effectiveness of control strategies. In future studies, incorporating this phenomenon will allow the model to make more realistic predictions and better quantify the robustness of control methods.

## Authors’ contributions

All authors contributed to the planning of the study, interpretation of results and editing of the manuscript. AP implemented the methods, ran the simulations and wrote the first draft of the manuscript.

## Acknowledgements

A large portion of this study was conducted during AP’s Master of Computer Science, which was jointly funded by the Faculty of Engineering and Information Technology at the University of Melbourne and Vingroup Science and Technology Scholarship Program.

We would like to thank Ben Phillips and Reid Tingley for their help in replicating their previous studies on exclusion, their valuable expertise on the behaviour and status of cane toads in Australia, as well as their feedback regarding the design of the models.

We acknowledge the Wurundjeri people, the Traditional Owners of the land on which the work described in this article was carried out, and pay our respects to their Elders past, present and emerging.

## Data Accessibility Statement

The data and code used in this study are publicly available on:

https://github.com/bda-pham/multiscale-canetoadcontrol.

