## Supporting Information for "Multiscale modelling of invasive species control: the cane toad in Australia"

---

In this Supporting Information document, we provide Overview–Design concepts–Details (ODD) descriptions of the two models as implemented. The ODD protocol was designed to promote clarity, rigour and reproducibility in descriptions of agent-based models (Grimm et al. 2006). In our descriptions, the components, parameters and assumptions of the models are explained in sufficient detail so that the models can be replicated. In addition, we provide the repository to access the code and data from the study as well as additional figures relevant to the manuscript.

#### S1 Microscale model - ODD description

##### S1.1 Overview

###### S1.1.1 Purpose

The purpose of the microscale model is to estimate the rate of spread of cane toads between a small number of water sources in one wet season and the impact of microscale processes, including deployment of control methods such as trapping and fencing, on this spread.

###### S1.1.2 Entities and state variables

In the model, there are two types of entity: mobile agents, such as toads, and immobile environmental features, such as water points, traps and fence sections. Toads are characterised by their gender, speed and location. Water points are characterised by their location, capacity and colonisation status (uncolonised, colonised and emitting). Traps are characterised by their location and capture count. Fence sections are characterised by their location and status (whether they are broken).

###### S1.1.3 Scales

The model is spatially explicit. The modelled space represents an abstract region between two arbitrary water points, mapped to a grid. The size of each grid-cell is 0.01 km$^2$ (100m x 100m) – this aligns well with the assumed water detection radius of toads (100m) (Tingley

Table S1: Attributes and variables of entities in microscale model

| Type of entity | Permanent attribute | State variable |
|---|---|---|
| Toad | gender, speed, heading deviation | location, heading |
| Water point | location, capacity | colonisation status |
| Trap | location | capture count |
| Fence section | location | status |

et al. 2012) and range of advertisement calls (120m) used as bait in traps (Muller et al. 2016, Muller & Schwarzkopf 2017) for optimising purpose. As locations of toads and water points are represented as real-valued coordinates, the grid size has no significant impact on the model except fence sections, which are modelled to be one grid-cell in length.

Simulations run in discrete time for 160 timesteps, each timestep indicating a day. The duration of 160 days serves as an upper bound of active days during wet seasons – from rainfall data, the highest number of active days is 127 (*Bureau of Meteorology* n.d.).

##### S1.1.4 Process overview

Here we provide a brief summary of how important processes are scheduled in the model. Details regarding those processes can be found in Section S1.3.3.

At every timestep:

1. Toads move if they are not at an uncolonised water point and not impeded by fence sections
2. Traps capture nearby toads with a probability
3. New water points are colonised if the colonisation conditions are met
4. Traps are reset if the timestep matches reset interval
5. All fence sections are fixed if the timestep matches fix interval

#### S1.2 Design concepts

The model revolves around the spread of a population – cane toads – through individual behaviours and the impact of environmental elements, such as water points and human-set traps, on this spread. Individual toads have basic objectives to survive and to breed, although these objectives are not always directly reflected in their behaviour. Toads can sense nearby water sources and move towards them, hence displaying adaptive behaviour; otherwise their movement is stochastic. Interactions between entities include implicit breeding between toads at water points, capture by traps, and fence sections impeding toad's movement. As a whole, the population of cane toads exhibits a collective behaviour of colonising new water sources at a certain probability. There is no learning in this model.

#### S1.3 Details

##### S1.3.1 Input data

The model requires no input at runtime.

##### S1.3.2 Initialisation

Here we provide details on how entities, model parameters and variables are initialised at the beginning of each simulation.

Water points are created using the model parameters wp-distribution and distance. Wp-distribution determines the number of water points in the model. By default it takes the value of "one-to-one", simulating the spread between two water points, one colonised and one uncolonised. When only modelling and studying toads' movement, it is set to "one-source". In "one-to-one" scenarios, the parameter "distance" determines the distance between the two water points.

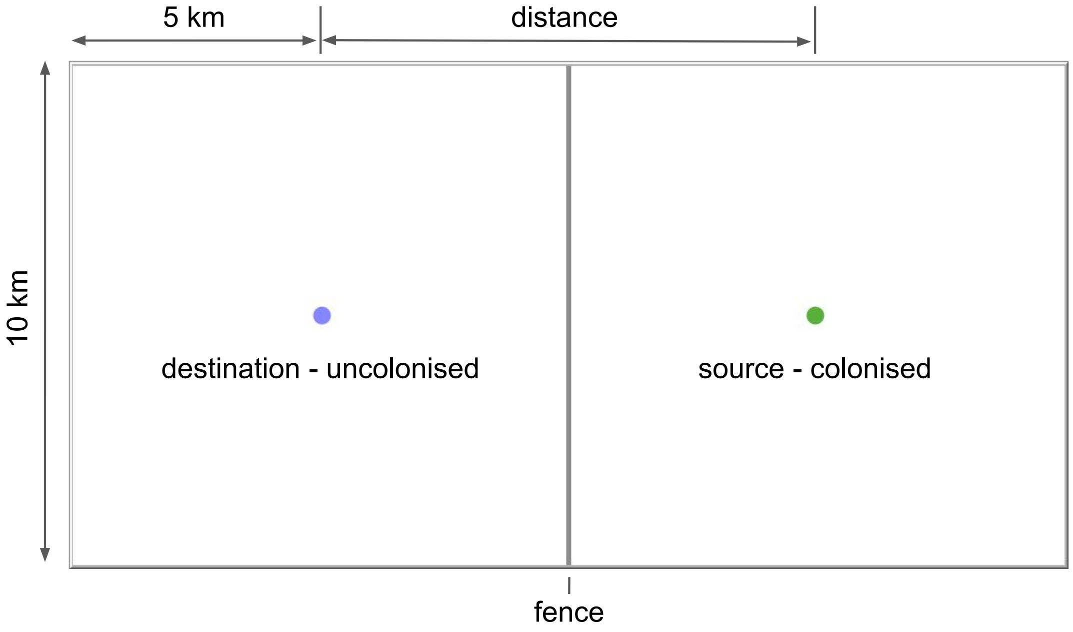


Figure S1: Model visualisation showing one-to-one scenario with a fence

Toads are created at emitting water points at every timestep. Each toad's initial heading is random (uniformly sampled between 0 to 360 degrees). The number of toads created is determined by toads-per-wave, which represents the emitting water point's capacity. Toads' gender is randomly initialised to Male or Female with a 1:1 ratio. Toads' speed is drawn from a normal distribution depending on their gender – this distribution is described in movement data from a field survey (Phillips et al. 2007). Toads' heading deviation, controlled by their attribute angle-dev, is drawn from a uniform distribution with the mean determined by mean-angle-dev. This value is calibrated to the recorded meander ratio in field survey

(Phillips et al. 2007). Finally, the distance at which toads are able to detect and move towards water points is assumed to be 100 m (Tingley et al. 2012).

Traps share the same attributes – there is no variation among traps other than their location and capture count. Their capture rate, range and capacity are taken from field studies (Muller & Schwarzkopf 2018, 2017, Muller et al. 2016). In the same studies, traps are often reset (i.e. removing captured toads) daily. Regarding deployment of traps, we assume they will be deployed in a square area of 200m x 200m centred on the water point and in density of 1 per 100m, meaning traps are placed 100m away from each other.

Fence sections are created in a straight line formation between two water points (Figure S7) and have one state variable representing whether they are damaged or not. We assume that fence sections have the same chance to be damaged (0.5% every day) and are all repaired periodically, with repair frequency determined by fence-fix-interval.

Table S2: Model parameters and initial value of state variables

| | | | |
|---|---|---|---|
| | | **Toads - global** | |
| male-speed-mean (m/day) | 46.2 | male-speed-dev | 31.3 |
| female-speed-mean (m/day) | 150.2 | female-speed-dev | 117.1 |
| mean-angle-dev (degrees) | 20 | water-detection-radius | 100 (m) |
| | | **Toads** | |
| gender | 50% male, 50% female | speed (m/day) | drawn from a normal distribution with speed-mean and speed-dev |
| location | at emitting water point(s) | angle-dev (degrees) | drawn from a uniform distribution with mean mean-angle-dev |
| heading (degrees) | random, 0 - 360 | | |
| | | **Water points - global** | |
| wp-distribution | one-to-one | distance (km) | as per scenario, 0 - 20 |
| toads-per-wave | 320 - 1600, default 960 | days-per-wave | 1 |
| | | **Water points** | |
| colonised? | 1 True, 1 False | emitting? | like colonised? |
| | | location | as per scenario, depending on distance |
| | | Traps - global | |
| trap-range | 120m | trap-reset-interval | 1 day |
| male-capture-rate (%/day) | 3 | female-capture-rate (%/day) | 3 |
| trap-density (traps/100m) | 1 (default) | trap-radius | 100m |
| max-capture | 30 | | |
| | | **Traps** | |
| Captures | 0 | Location | laid out in uniform distribution around water points, from trap-density and trap-radius |
| | | **Fence - global** | |
| fence-break-prob (%/day) | 0.5 | fence-fix-interval (days) | 7 (default) |
| | | **Fence sections** | |
| broken? | false | location | form a line in the middle, between 2 water points |

##### S1.3.3 Submodels

Here we provide details regarding processes within the model. This includes:

- Toad movement – how toads move at each timestep and how this movement changes with the presence of nearby water points and fence sections
- water point colonisation – how colonisation status of water points changes
- Trap – the procedure of trap captures and capacity
- Fence – how the broken status of fence sections change

**Toad movement** At every timestep,

- if there is an uncolonised water point within water-detection-radius, move towards it a distance determined by speed; otherwise
- heading is drawn from a normal distribution with previous heading as the mean and toad-angle-dev as the deviation, then
- move a distance specified by the toad's individual speed towards heading. If the path is blocked by a fence section, stop where the fence section is. If the path crosses an uncolonised water point within water-detection-radius, change heading towards it.

**Water point colonisation** At every time step, each uncolonised water point performs a check. If there are at least one toad from each gender at the water point, it is colonised. Once a water point is colonised, it remains colonised. If there is no more uncolonised water point, the model stops.

**Trap** At every timestep, if a toad is within the range of a trap, it has a chance to be captured. Captured toads are removed from the model, and the capture count of the trap is increased by one for each toad captured. Each trap has a capacity, and can no longer capture toads if captures reach capacity. The capture count of traps is reset periodically.

**Fence** As mentioned in Toad movement submodel, toads cannot move past a fence section. At every timestep, each section of the fence has a chance to be damaged (fence-break-prob). Toads can cross damaged sections freely. The entire length of the fence is repaired periodically with the repair frequency determined by fence-fix-interval.

### S2 Macroscale model - ODD description

#### S2.1 Overview

##### S2.1.1 Purpose

The purpose of the macroscale model is to estimate the spread of cane toads in the Kimberley-Pilbara corridor over many years and the impact of control strategies (the deployment of one or more control methods in an area) on this spread.

##### S2.1.2 Entities and State variables

In the model, there is only type of entity: immobile environmental features, such as water points and weather stations. Water points are characterised by their location, size, capacity, colonisation status and control status. Weather stations are characterised by their location, the current number of active days and the statistics describing the distribution of active days at that station (mean, max, min and standard deviation).

Table S3: Attributes and variables of entities in macroscale model

| **Type of entity** | **Permanent attribute** | **State variable** |
|---|---|---|
| Water point | location, size, capacity, trapped?, fenced?, excluded? | colonised?, emitting?, exclusion-fail?, exclusion-fail-since |
| Weather station | location, mean-active-days, max-active-days, min-active-days, dev-active-days | active-days |

##### S2.1.3 Scales

The model is spatially explicit. The modelled space represents the Kimberley-Pilbara corridor, approximately 400 km x 300 km in size. Simulations run in discrete time for at most 200 timesteps, each timestep indicating a year.

##### S2.1.4 Process overview

Here we provide a brief summary of how important processes are scheduled in the model. Details regarding those processes can be found in Section S2.3.3.

At every timestep:

1. Weather stations generate active-days for the current timestep
2. Working exclusion mechanisms at water points have a chance to fail

3. Failed exclusion mechanisms get repaired after 2 timesteps
4. Water points colonised in previous timestep start to emit colonisers
5. Uncolonised water points are colonised with a probability by nearby emitting water points

#### S2.2 Design concepts

The model revolves around the spread of cane toads by colonising water points in a region, the rainfall pattern which influences this spread, and control strategies aimed to deter this spread. Water points and weather stations are static, passive entities and do not have objectives. Interactions between entities include colonised water points attempting to colonise uncolonised water points, and weather stations determining rainfall patterns of nearby water points. As a whole, the collection of water points behaves like a network through which toads spread across the landscape. There is no sentient agent and thus no adaptation or learning in this model.

#### S2.3 Details

##### S2.3.1 Input

The model requires no input at runtime.

##### S2.3.2 Initialisation

Here we provide details on how entities, model parameters and variables are initialised at the beginning of each simulation.

Water points are created from water point data, which contain the coordinates and type of each water point. If the water is an irrigation area, its capacity is set as the product of mean-capacity and its area in square kilometres. Otherwise, capacity is drawn from a Poisson distribution with a mean determined by mean-capacity. All variables concerning control are set to False. Most water points are set to be uncolonised and not emitting, except those in North-Eastern corner, which are set to colonised and emitting to simulate the invasion having reached the corridor. Links are created between water points that are close enough to colonise each other.

Weather stations are created from the aggregated water station data (*Bureau of Meteorology* n.d.), which contain their location and statistics regarding the distribution of active days. The interval between wet years is set to 10 years by default.

Tables containing colonisation probability of water points are loaded from data (Figure S3). These include one table of base spread without control and three tables of spread under different controlled scenarios (fencing; trapping; fencing and trapping).

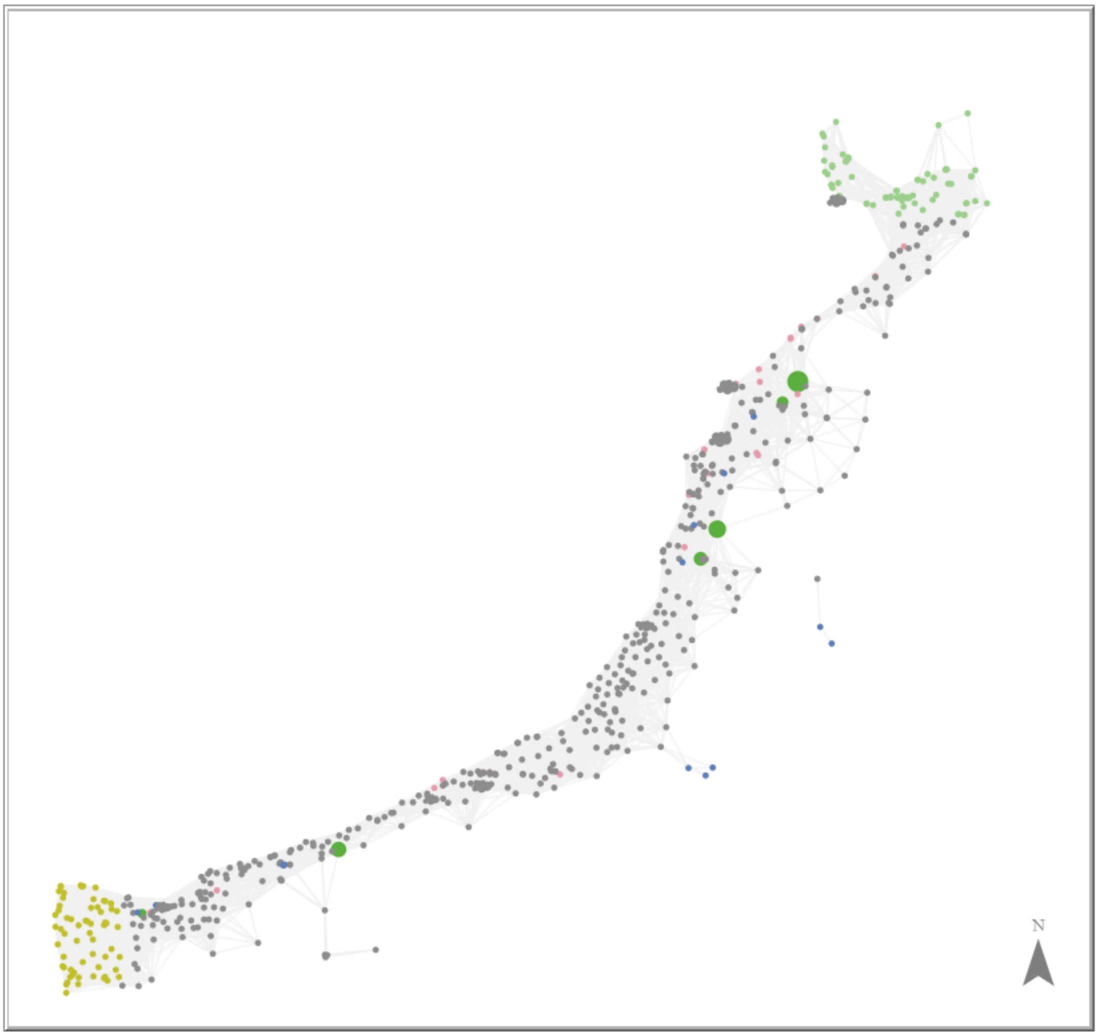


Figure S2: Model visualisation showing water points in the Kimberley-Pilbara corridor. The points represent water points, with the colour representing the type of each water point – blue for natural water points, red for dwellings, dark green for irrigation areas and light grey for other types of artificial water points. Water points are linked to nearby water points within colonisation distance. Grey cloud shapes are weather stations. At the beginning of the model, water points in the North-Eastern corners are initialised as colonised, marked visually by their light green colour. If the toads reach any of the water points in the South-Western corner, marked by their yellow colour, the region is considered to be completely colonised and the simulation ends.

From a series of pre-generated locations evenly spaced along the corridor, at most one control location is active during each simulation. A number of water points closest to this location are identified (no-controlled-wps), and control methods are enabled at those water points (excluded? and trapped?). If fencing is enabled, mark links (i.e. colonisation trajectories) that cross the corridor-fence (fenced? = true).

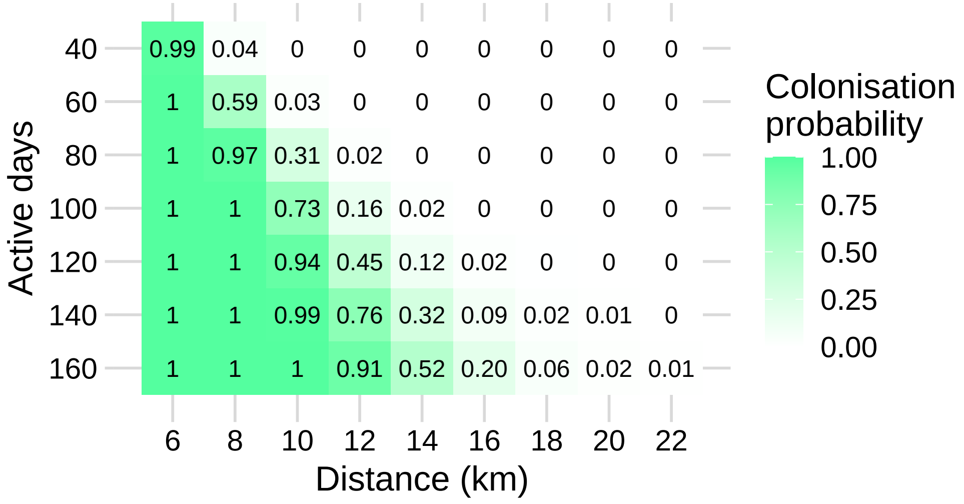


Figure S3: Function of colonisation probability as interface between models. Only two variables, distance and active days, are present in this figure. The values shown in the figure are from a capacity of 1600 colonisers per day.

Table S4: Model parameters and initial value of state variables

| | | | |
|---|---|---|---|
| | **Water points - global** | | |
| mean-capacity | 960 | exclusion-fail-prob | 0.05 |
| exclusion-repair-delay (years) | 2 | | |
| | **Water points** | | |
| location and type | from data | capacity | drawn from a Poisson distribution with mean mean-capacity |
| colonised? and emitting? | false | trapped? and excluded? | false |
| exclusion-fail? | false | exclusion-fail-since | -1 |
| | **Water point links** | | |
| fenced? | false | | |
| | **Weather stations - global** | | |
| wet-year-interval (years) | 10 | wet-year-extra-days | 14 |
| | **Weather stations** | | |
| location, mean-active-days, max-active-days, min-active-days, stddev-active-days | from data | active-days | 0 |
| | **Control - global** | | |
| no-controlled-wps | as per scenario | location | from data |
| active-control-loc | 0 - 16, as per scenario | exclusion? | as per scenario |
| trap | none / density 1 / density 2, as per scenario | fence | none / weekly repair / monthly repair, as per scenario |

##### S2.3.3 Submodels

Here we provide details regarding processes within the model, including:

- Active days: how the number of active days is generated and assigned to water points

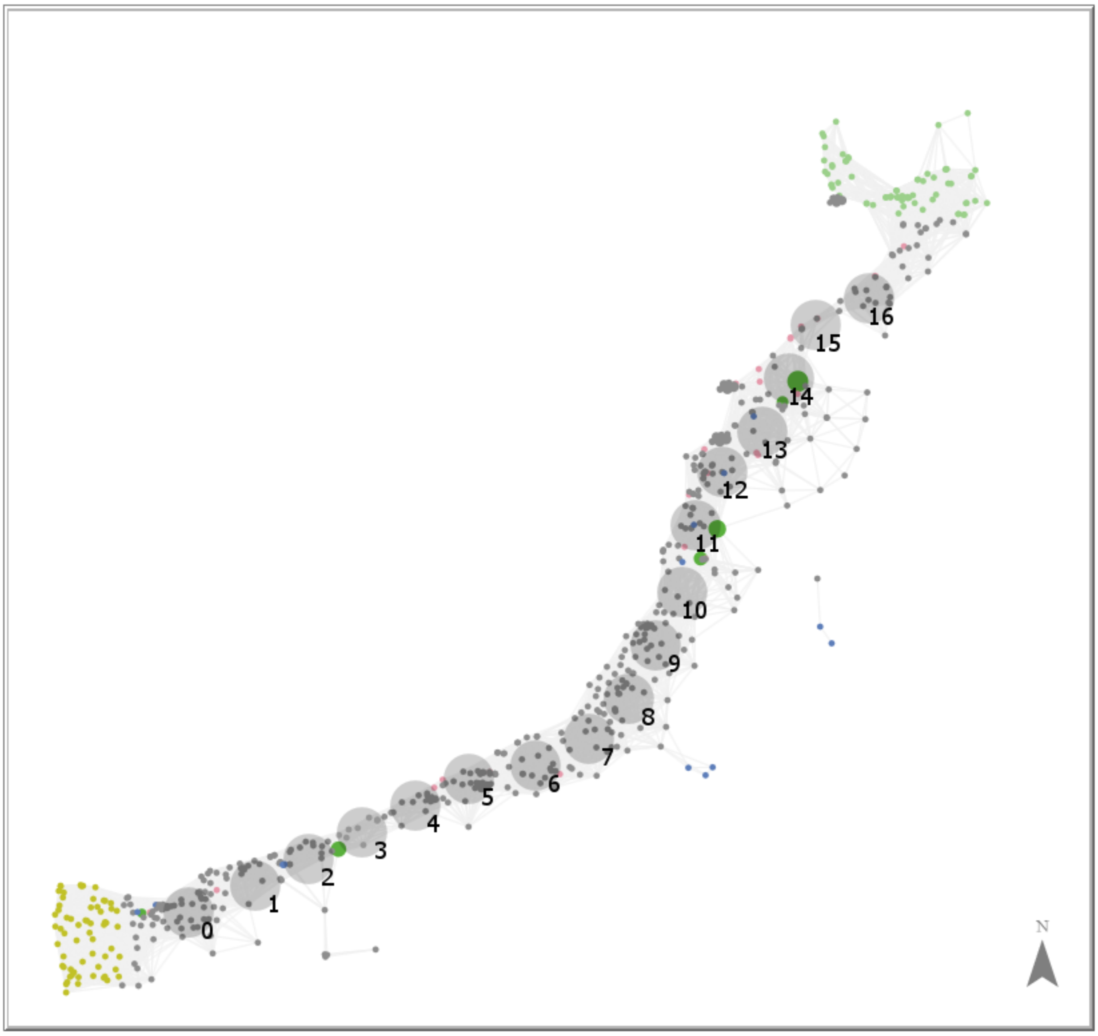


Figure S4: Control locations are evenly positioned along the corridor

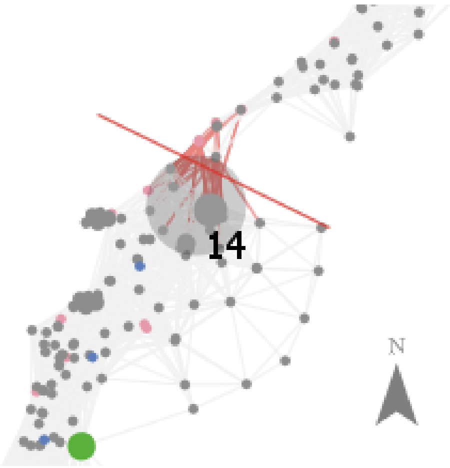


Figure S5: Corridor-fence affects all links that intersect it

- Colonisation probability: how colonisation probability is looked up and modified
- High capacity: how capacity above the limit is modelled by modifying the look-up distance
- Exclusion: how exclusion status of water points changes
- Trap and fence: how traps and the corridor-fence affect the colonisation process

**Weather stations and active days** At every timestep, the number of active days at a weather station is generated from a normal distribution described by its statistics (mean-active-days, max-active-days, min-active-days, stddev-active-days). However, if the current timestep falls on a wet year, a weather station's active-days is set at max-active-days + wet-year-extra-days instead. The number of active days (days in which toads are able to move) at a waterbody is modelled to be the same as the closest weather station. The number of active days of a colonisation trajectory – a directed edge between two water points – is equal to the number of active days at the destination.

**Colonisation probability** At every timestep, for each uncolonised water point, emitting water points in range attempt to colonise it one by one. Each source's probability of colonising a new water point depends on the source's capacity, the distance between them, as well as the number of active days of the uncolonised water point. Those three parameters determine where in the spread table the colonisation probability should be looked up. As the actual parameters in each case might fall between the values used to generate the spread table, the colonisation probability for a set of parameters (capacity, distance, active-days) is computed using trilinear interpolation (Bourke 1999), a common approach to interpolate between 8 points in a 3-dimensional space. If a parameter is smaller than the lower bound of their range in the spread table, it is treated as the lowest (since the lowest distance results in near-certain colonisation anyway). For example, water points less than 6 km away from each other have the same colonisation probability as if they are 6 km (lowest distance) away from each other.

If the water point has an area measurement, the area is assumed to be circular and the radius is computed accordingly. Afterwards, the probability of such a water point is augmented following the formula:

$$Pr_{colon}(..., radius) = Pr_{colon}(...) \cdot (1 + (radius * 10/4)) \quad (1)$$

This formula is based on the correlation between water detection radius and colonisation probability in the microscale model (Figure S8). Given the small number of water points with area measurement, we assume this linear approximation is adequate.

**Capacity of water points** Capacity of each water is drawn from a Poisson distribution with mean mean-capacity. For water points of type irrigation area, their capacity $C_i$ is calculated following the formula in (Southwell et al. 2017). Specifically:

$$C_i = C \times A_i \times D \tag{2}$$

where $C$ is the mean capacity, $A_i$ is the area of the irrigated area, and $D$ is the mean density of water points in the corridor. The mean density $D$ was calculated in (Southwell et al. 2017) by dividing the number of water points (566) by the area of the Kimberley-Pilbara corridor (15,402 km$^2$).

To approximate capacities that exceed the highest capacity in the spread table, when looking up colonisation probability in the spread table we compute an equivalent combination of capacity and distance. Specifically, the source's capacity is divided by 4 while distance is reduced by 2 km. This approximation is applied until capacity falls within the range of (20 - 100), and the resulting capacity and distance can then be used to look up the colonisation probability. This procedure is based on the relationship between capacity and distance (Figure S9), specifically how a multiplication of capacity by 4 allows toads to spread approximately 2km further with comparable probability. Formally:

$$Pr_{colon}(4 \cdot capacity, distance, ...) = Pr_{colon}(capacity, distance - 2, ...) \tag{3}$$

**Exclusion** Water points that are controlled with exclusion (excluded? is True) cannot be colonised. At every timestep, each of those water points has a chance (exclusion-failure-prob) to become available for colonisation (set exclusion-fail? to True). When this failure happens, the timestep is recorded (exclusion-fail-since), and after a number of timesteps (exclusion-repair-delay), the failure is reverted (set colonised?, emitting? and exclusion-fail? to False).

**Trap and fence** When a water point is attempting to colonise another, if the uncolonised water point is controlled with traps or the link between them is fenced, the colonisation probability is looked up in the corresponding spread table. The spread table is chosen based on the control status of the destination water point and the trajectory between the two water points following the rules described in Table S5.

Table S5: Spread table from control statuses of trajectory and destination

| Trajectory - fenced? | Destination - trapped? | Spread table |
|---|---|---|
| true | true | fence-trap |
| true | false | fence |
| false | true | trap |
| false | false | base |

### S3 Code and data

The NetLogo source code of the models, input data for the macroscale model, and R scripts used for data processing and result analysis can be found in this repository: `https://`

github.com/bda-pham/multiscale-canetoadcontrol.


#### S4 Supplementary Figures

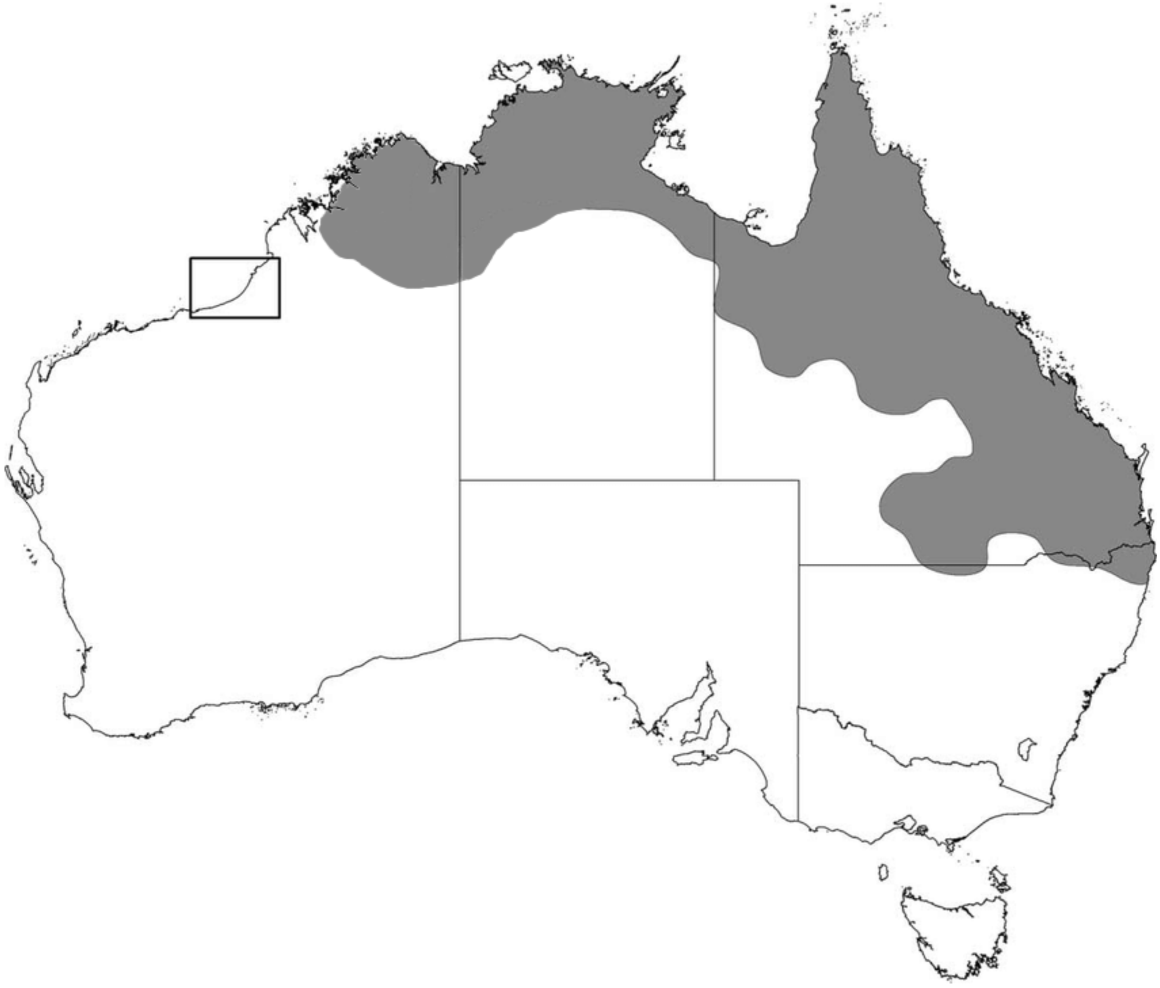

Figure S6: Approximate distribution of Cane toads in Australia as of 2022. The small box represents the Kimberley-Pilbara corridor in Western Australia.

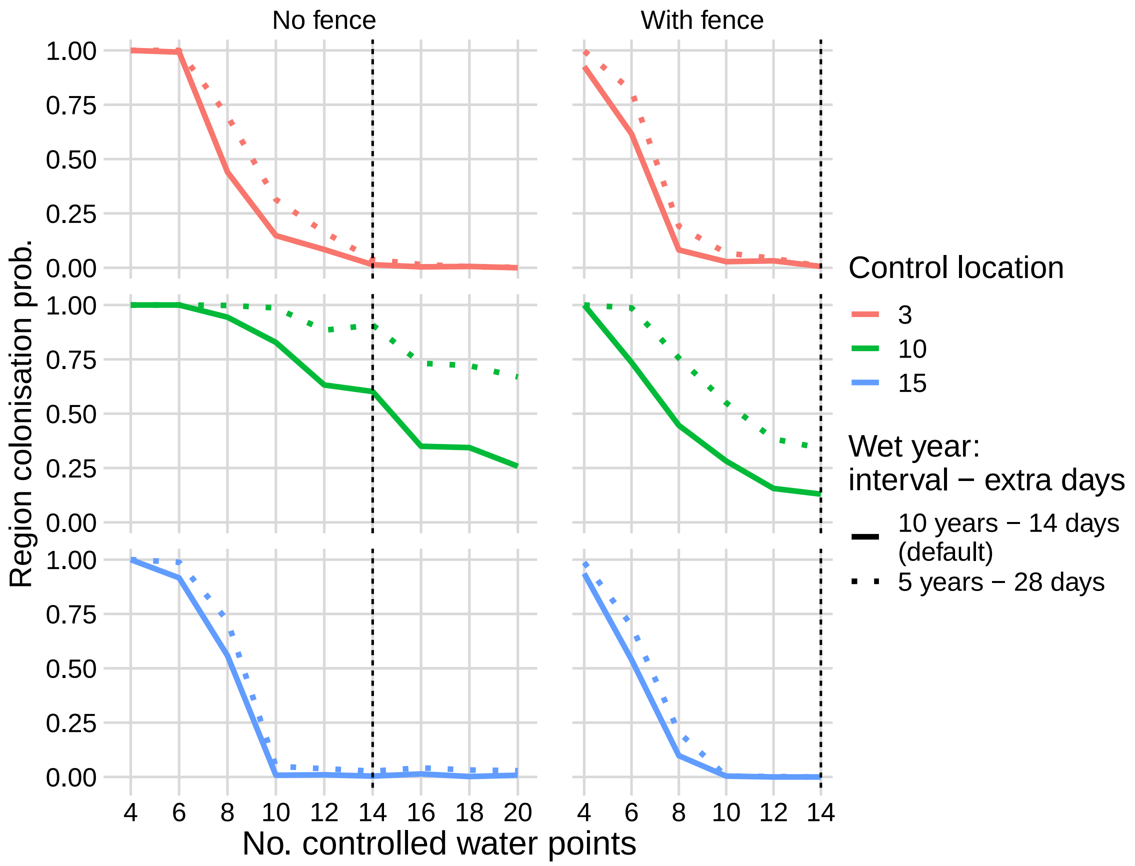


Figure S7: Wetter climate slightly reduces the effectiveness of control methods.

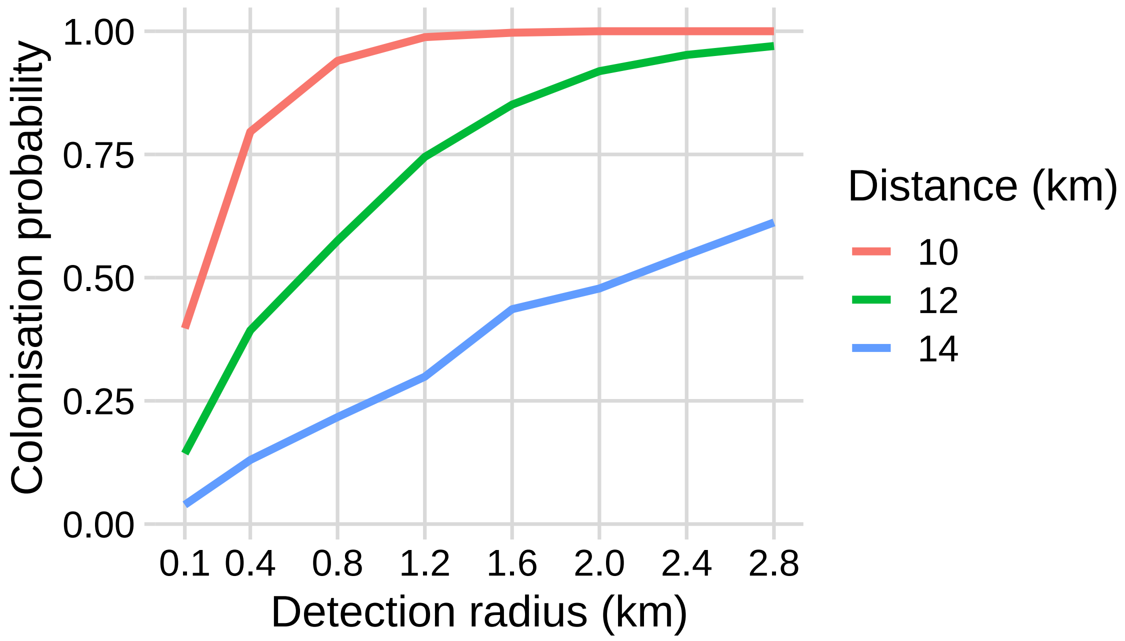


Figure S8: Correlation between detection radius and colonisation probability.

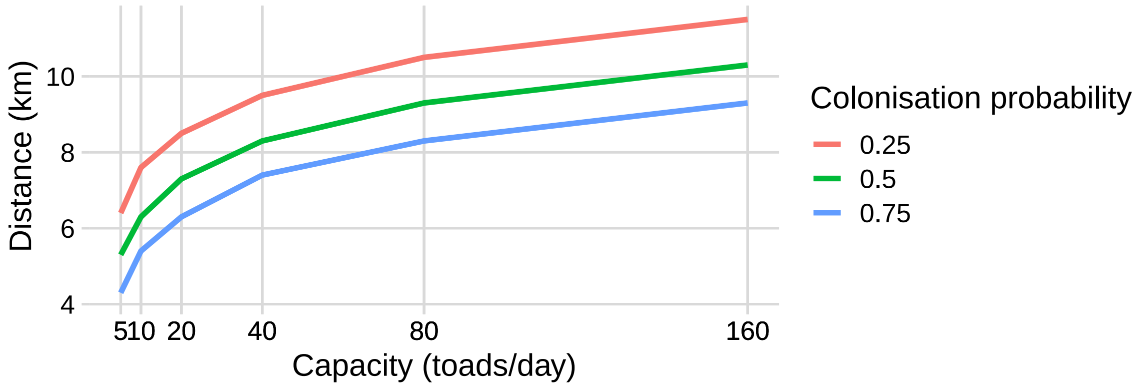


Figure S9: Correlation between detection radius and colonisation probability.
